# Genome-wide transcriptional responses of marine nematode *Litoditis marina* to hyposaline and hypersaline stresses

**DOI:** 10.1101/2021.02.21.432187

**Authors:** Yusu Xie, Pengchi Zhang, Liusuo Zhang

## Abstract

Maintenance of osmotic homeostasis is essential for all organisms, especially for marine animals in the ocean with 30‰ salinity or higher. However, the underlying molecular mechanisms that how marine animals adapt to high salinity environment compared to their terrestrial relatives, remain elusive. Here, we investigated marine animal’s genome-wide transcriptional responses to salinity stresses using an emerging marine nematode model *Litoditis marina*. We found that the transthyretin-like family genes were significantly increased in both hyposaline and hypersaline conditions, while multiple neurotransmitter receptor and ion transporter genes were down-regulated in both conditions, suggesting the existence of conserved strategies for response to stressful salinity environments in *L. marina*. Unsaturated fatty acids biosynthesis related genes, neuronal related tubulins and intraflagellar transport genes were specifically up-regulated in hyposaline treated worms, while exhibited the opposite regulation in hypersaline condition. By contrast, cuticle related collagen genes were enriched and up-regulated for hypersaline response, interestingly, the expression of these collagen genes was significantly decreased in hyposaline condition. Given a wide range of salinity tolerance of the marine nematodes, this study and further genetic analysis of key gene(s) of osmoregulation in *L. marina* will likely provide important insights into biological evolution and environmental adaptation mechanisms in nematodes and other invertebrate animals in general.

## 1 Introduction

Salinity, as an important ecological factor, affects the physiology and behavior of marine and terrestrial animals. As a nutrient element in the diet, salt is of vital importance to the health of animal and human. In human, chronic high dietary salt intake gradually causes an increased risk for cardiovascular disease, particularly hypertension; as well as other disease such as stroke, gastric cancer, kidney disease and obesity (Rust and Ekmekcioglu, 2017;He and MacGregor, 2018). Therefore, studies on the underlying mechanisms of animals’ sensation, response, and adaptation to environmental salinity have always been a hot topic.

The multicellular model organism, *Caenorhabditis elegans* senses most of the environmental disturbance by the terminal cilia of sensory neurons (Bargmann, 2006). It is known that ASH neurons are required for the perception of high salt, while ASEs are for the low salt. The sensation of salinity stress can trigger subsequent avoidance behavior to protect worms from harmful salinity conditions (Hilliard et al., 2005;Kunitomo et al., 2013). Once the stressed salinity environment is unavoidable, *C. elegans* will engage a sleep-like quiescent behavior and cease locomotion and feeding, which is dependent on ALA neuron (Hill et al., 2014). Due to the imbalance between internal and external osmotic pressure, the body volume of nematodes undergoes significant changes under salinity stresses, manifests as shrinkage under hypersaline, and swelling under hyposaline (Lamitina et al., 2004). Organic osmolytes play an important role in osmotic regulation and salinity stress adaptation for all organisms. In *C. elegans*, cellular osmotic homeostasis can be maintained by rapid accumulation of glycerol upon high salinity challenge (Lamitina et al., 2004;Lamitina et al., 2006). It is well accepted that the *C. elegans*’ cuticle might act as a “sensor” in responding to salinity stress damage, which in turn triggers downstream physiological changes (Choe, 2013;Dodd et al., 2018). On the other hand, numerous genes involved in osmotic regulation have been identified in *C. elegans*, such as the osmolyte glycerol synthesis enzyme gene (Lamitina et al., 2004;Lamitina et al., 2006;Choe, 2013), transient receptor potential cation channel TRP subfamily genes (Choe, 2013), chloride channel genes (Choe, 2013), aquaporin water channel genes (Igual Gil et al., 2017), extracellular matrix component genes (Lamitina et al., 2006;Choe, 2013), as well as genes related to MAPK, WNK-1/GCK-3, Notch and insulin-like signaling pathways (Choe, 2013;Dresen et al., 2015;Burton et al., 2017). Many of the above osmotic regulation genes play evolutionarily conserved roles in systemic osmotic homeostasis in yeast, flies, plants and mammals (Strange et al., 2006;Burg et al., 2007;Brewster and Gustin, 2014;Pasantes-Morales, 2016;Zhou et al., 2016;Yang and Guo, 2018), providing clues for treatment of human disease that accompany osmotic perturbation.

*C. elegans* is one of the typical free-living terrestrial nematode species, whereas about 43% of the known nematode species are distributed in the ocean (Appeltans et al., 2012;Zhang et al., 2015). It is speculated that nematodes may have emerged from a marine habitat during the Cambrian explosion (van den Elsen et al., 2009), and colonized land about 442 million years ago (Rota-Stabelli et al., 2013). Salinity is obviously one of the most significant factors that changed during this successful terrestrialization. However, the underlying mechanisms are largely unexplored.

*Litoditis marina* is a dioecious free-living marine nematode, which is widely distributed in the littoral zone of coasts and estuaries, and plays an important role in these marine ecosystems (Derycke et al., 2016;Xie et al., 2020). It possesses some promising characteristics similar as *C. elegans*, such as short generation time, clear genetic background and a sequenced genome (Xie et al., 2020), which facilitated its laboratory application for the in-depth study of molecular biology, cell biology, physiology and behavior regulation in this species. Generally, the habitat salinity for intertidal marine nematodes, including *L. marina*, is frequently changed due to the influence of many factors such as tides, sun exposure, rainfall, ocean currents and climate. The effective sensation and response to the dynamic salinity environments is of great significance for marine nematodes’ survival. However, the underlying molecular mechanism is still unknown.

In this study, we challenged *L. marina* L1 larvae with hyposaline and hypersaline stresses respectively, and further demonstrated their genome-wide transcriptional signatures via RNA sequencing (RNA-seq) analysis. Both common and specific responding genes were identified in hyposaline and hypersaline stressed worms. These results not only provide a basis for understanding the salinity response mechanism for *L. marina*, but also might provide new clues for in-depth exploration of osmoregulation and environmental adaptation mechanisms for other marine animals.

## 2 Materials and Methods

### 2.1 Worms

The wild strain of marine nematode *L. marina*, HQ1, was isolated from intertidal sediments (Huiquan Bay, Qingdao). Healthy worms were cultured on SW-NGM agar plates (prepared with seawater with a salinity of 30‰) seeded with a lawn of *Escherichia coli* OP50 as a food source, as reported previously (Xie et al., 2020). Worms were maintained and propagated at 20°C in the laboratory for about 3 years till this study.

### 2.2 Behavioral and Developmental Analysis under Salinity Stresses

Three sets of salinity conditions were applied for salinity-stress treatment. Artificial seawater-NGM agar plates were prepared by Sea Salt (Instant Ocean) in 3‰ (hyposaline), 30‰ (control) and 60‰ (hypersaline) salinity, respectively.

For behavioral and developmental analysis, 30 newly hatched L1s were transferred onto each indicated 3 cm-dimeter agar plates seeded with 15 μl OP50. Worms were scored as active if response was detected after prodding with a platinum wire 24 h post-treatment. The number of adult worms was scored 120 h (5 days) post-treatment. Three replicates were performed for each experimental condition.

### 2.3 RNA-seq Analysis

HQ1 strain worms cultured on SW-NGM plates were allowed to lay eggs overnight at 20°C. Eggs were washed off and collected using filtered sterile seawater, then treated with Worm Bleaching Solution (Sodium hypochlorite solution : 10 M NaOH : H_2_O = 4 : 1 : 10, prepared in terms of volume ratio) at room temperature for 1.5 min. Wash eggs twice with sterile seawater. Leave the worms to hatch overnight and undergo growth arrest in sterile seawater at 20°C. Synchronized L1 worms were collected by filtration using 500 grid nylon filter of 25 μm mesh size, and then transferred to each 9 cm-dimeter agar plates prepared by Sea Salt mentioned above, which were seeded with 100 μl OP50 per plate (covering the entire plate evenly with a coating stick), respectively. Treated L1s were collected after incubating for 3 h at 20°C under each salinity condition. Worms were washed with M9 for three times to remove the bulk of the residual bacteria. Excess supernatants were removed carefully via centrifugation. The samples were frozen immediately in liquid nitrogen. Total RNA was then extracted using Trizol (Invitrogen).

With three biological replicates for each treatment, a total of nine RNA libraries were prepared with 3 μg RNA using NEBNext® UltraTM RNA Library Prep Kit for Illumina® (NEB, USA) following manufacturer’s recommendations. Then, RNA libraries were sequenced on an Illumina NovaSeq 6000 platform and 150 bp paired-end reads were generated.

Clean data, with Q20 value higher than 97.5 for each sample, were first obtained by removing reads containing sequencing adaptors, reads having poly-N and low-quality ones from raw data. Then, they were aligned to the *L. marina* reference genome (Xie et al., 2020) by Hisat2 (v2.0.5) (Kim et al., 2015). New transcripts for novel genes were predicted and assembled by StringTie (v1.3.3b) (Pertea et al., 2015), then annotated with Pfam, SUPERFAMILY, GO and KEGG databases (Kanehisa and Goto, 2000;Young et al., 2010). Further, the reads numbers mapped to each gene were analyzed using featureCounts (v1.5.0-p3, with parameter −Q 10 −B −C) (Liao et al., 2014), and FPKM (expected number of Fragments Per Kilobase of transcript sequence per Millions base pairs sequenced of each gene) was calculated based on the length of the gene and reads count mapped to this gene, which was used for estimating gene expression levels. Differential expression analysis of two conditions was performed using the DESeq2 R package (v1.16.1) (Love et al., 2014). The resulting *P*-values were adjusted using the Benjamini and Hochberg’s approach for controlling the false discovery rate. Genes with an adjusted *P*-value < 0.05 found by DESeq2 were assigned as differentially expressed. Moreover, we used clusterProfiler R package (v3.4.4) to test the statistical enrichment of differential expression genes in Gene Ontology (GO) terms and KEGG pathways, the corrected *P*-value < 0.05 were considered significantly enriched by differential expressed genes.

### 2.4 Real-Time PCR Analysis

Some of the key genes of our interest were selected for qPCR validation: transthyretin-like family gene EVM0003534, trehalose-6-phosphate synthase gene EVM0007411/*tps-2*, dopamine receptor gene EVM0000190/*dop-1*, glutamate receptor gene EVM0013383/*glc-4*, acetylcholine receptor gene EVM0009741/*eat-2*, serotonin receptor gene EVM0012843/*ser-1*, neuropeptide receptor genes EVM0015448/*npr-6* and EVM0010018/*npr-4*, ion transporter genes EVM0004010/*kcc-2* and EVM0012374/*twk-24*, fatty acid elongation gene EVM0013022/*elo-2* and fatty acid desaturase gene EVM0001302/*fat-4*, tubulin gene EVM0007116/*tba-5*, cuticle collagen genes EVM0002243/*col-156* and EVM0005554/*col-107*.

Synchronized L1 worms were separately treated under each salinity condition (3‰, 30‰ and 60‰) using artificial seawater-NGM plates (prepared by Sea Salt, Instant Ocean) at 20°C for 3 h. Each treatment was performed for three biological repeats. Total RNA was extracted using Trizol (Invitrogen), reserve transcribed to cDNA using the ReverTra Ace® qPCR RT Master Mix with gDNA Remover kit (TOYOBO, Code No. FSQ-301), and the cDNA was used for qPCR analysis using the QuantStudioTM 6 Flex Real-Time PCR System (Applied Biosystems) and SYBR Green detection system (TOYOBO, Code No. QPK-201). The primers information of totally 15 salinity related genes, listed above, was shown in Supplementary file 1. Each experiment was performed in triplicates for each biological replica. Values were normalized against the reference gene EVM0013809, which is orthologue of *C. elegans* gene *cdc-42* (Hoogewijs et al., 2008). Gene expression was presented as a fold change using the delta Ct method (Livak and Schmittgen, 2001). Data were statistically analyzed by one-way analysis of variance (one-way ANOVA) using SPSS software 11.0; values were considered to be significant at *P* < 0.05.

## 3 Results

### 3.1 *L. marina* behavioral and Developmental Defects under Salinity Stresses

*L. marina* is maintained under 30‰ salinity condition in the laboratory, around 91% newly hatched L1 larvae developed into adulthood after 5 days at 20°C (**Figure 1B**). To test its salinity tolerance, we first treated L1 worms under two conditions: hyposaline with a 3‰ salinity and hypersaline with a 60‰ salinity. We observed that L1 worms were paralyzed immediately on both salinity plates, with obvious body volume change in a manner similar to that reported in *C. elegans* (Lamitina et al., 2004). Worms can recover their motility afterwards. Compared to the control group (30‰ salinity), approximately 88% worms under hyposaline could move normally after 24 h, whereas only 29.4% L1s could recover motility under hypersaline (**Figure 1A**). Next, we did the same test applied to even higher salinity conditions such as 70‰ and 80‰, and observed that worms cannot survive under those conditions, indicating 60‰ is the extreme high salinity for *L. marina* to tolerate.

**FIGURE 1.**
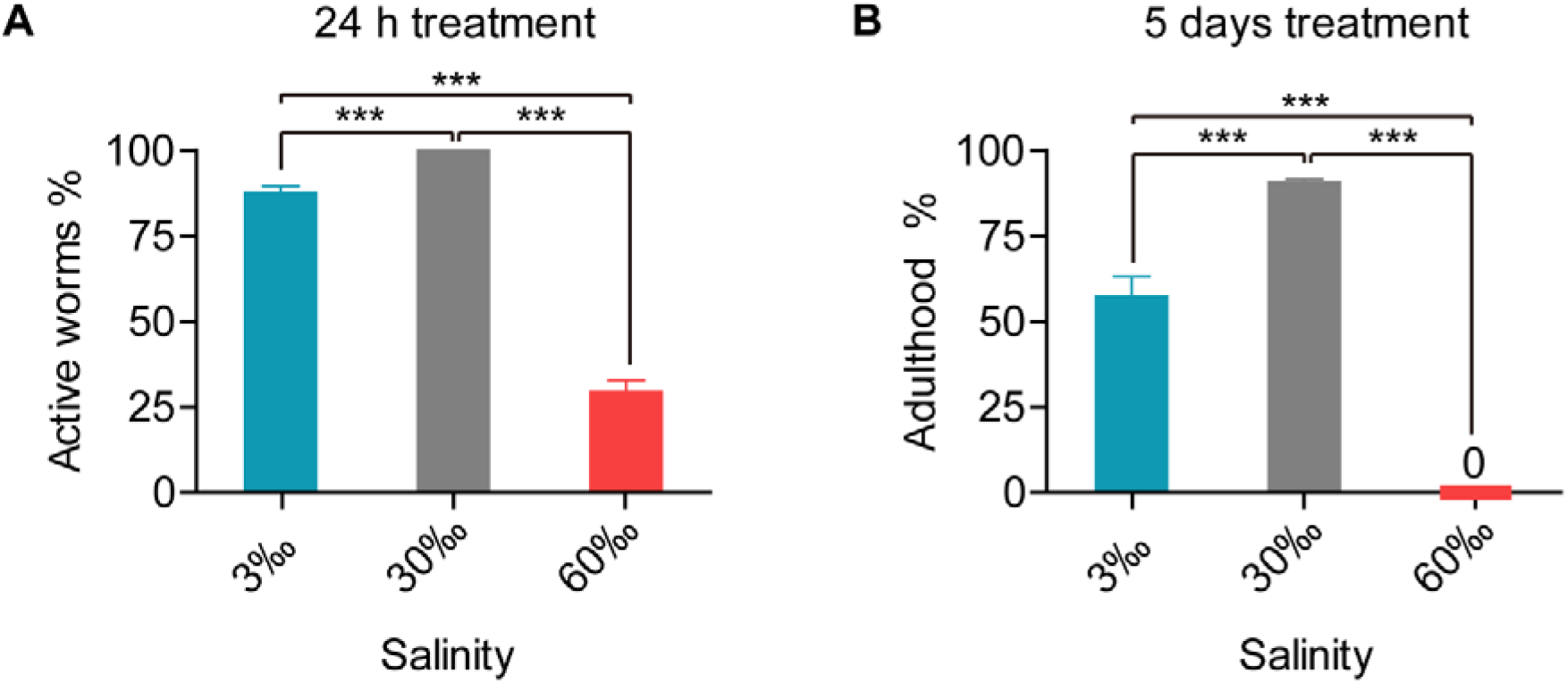
Behavioral and developmental defects in *L. marina* responding to salinity stresses. Marine nematode *L. marina* which is normally maintained under 30‰ salinity condition, showed both behavioral **(A)** and developmental **(B)** defects when stressed with either low salinity (3‰) or high salinity (60‰). Notably, worms showed enhanced defects under 60‰ salinity.

We further found that, upon the 5th day of treatment, 57.5% worms reached adulthood under hyposaline (3‰), while no adults was observed under hypersaline (60‰) condition. Thus, both hyposaline and hypersaline attenuated worms’ development (**Figure 1B**).

Taken together, worms exhibited both significantly behavioral and developmental defects when stressed with either low salinity or high salinity.

### 3.2 RNA-seq Analysis in *L. marina* under Hyposaline and Hypersaline Environments

To investigate genome-wide responses in *L. marina* to salinity stress, we used RNA-seq analysis. Newly hatched L1s were treated for 3 h on low salinity (3‰), normal salinity (30‰, control) and high salinity (60‰) plates, respectively (**Figure 2A**). A total of 1209 differentially expressed genes (DEGs) were identified under low salinity, and 1330 DEGs under high salinity. Interestingly, there were 108 up-regulated DEGs and 93 down-regulated DEGs shared in both conditions (**Figure 2A**), indicating common response patterns under hyposaline and hypersaline stresses. On the other hand, condition-specific DEGs exhibited salinity-dependent responsive and regulatory mechanisms in *L. marina*. Details of significantly up-regulated and down-regulated DEGs were listed in Supplementary file 2.

**FIGURE 2.**
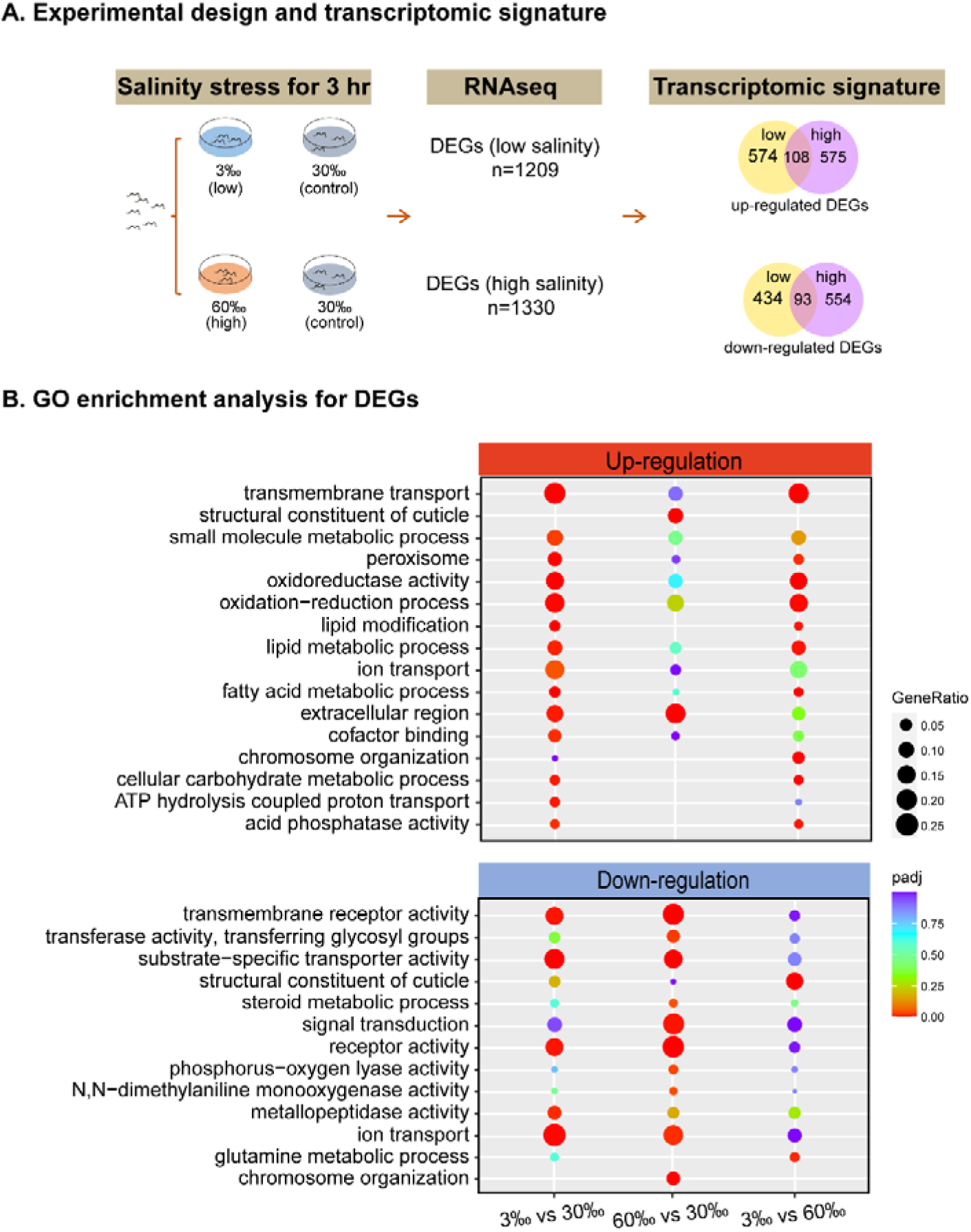
RNA-seq identifies the transcriptomic signature of short-time salinity stressed marine nematodes. **(A)** Experimental design of this study and the resulting transcriptomic signature of salinity stressed worms. Differentially expressed genes (DEGs, |log2foldchange|>1; DESeq2 padj<0.05) were determined for each condition. **(B)** GO enrichment analysis for DEGs. |log2foldchange|>1; DESeq2 padj<0.05 was set as the differential gene screening threshold.

Based on GO enrichment analysis for DEGs, we observed that there were more up-regulated GO terms significantly enriched under low salinity, whereas more down-regulated GO terms were significantly enriched under high salinity (**Figure 2B**). Specifically, extracellular region genes were up-regulated while receptor and transporter genes were down-regulated under both conditions (**Figure 2B**).

### 3.3 Shared Transcriptomic Signature of *L. marina* under both Low and High Salinity Stress Conditions

As both hyposaline and hypersaline stresses lead to behavioral and developmental defects in *L. marina*, common transcriptomic signature was found between these two conditions based on GO enrichment analysis.

As shown in Figure 2B, extracellular region related genes were significantly enriched in DEGs in both examined salinity conditions. We found that a series of transthyretin-like family genes, such as EVM0004638/*ttr-30* and EVM0003584/*ttr-48*, were up-regulated under both conditions (**Figure 3A**), indicating that extracellular region related genes can be induced by either low or high salinity stress.

**FIGURE 3.**
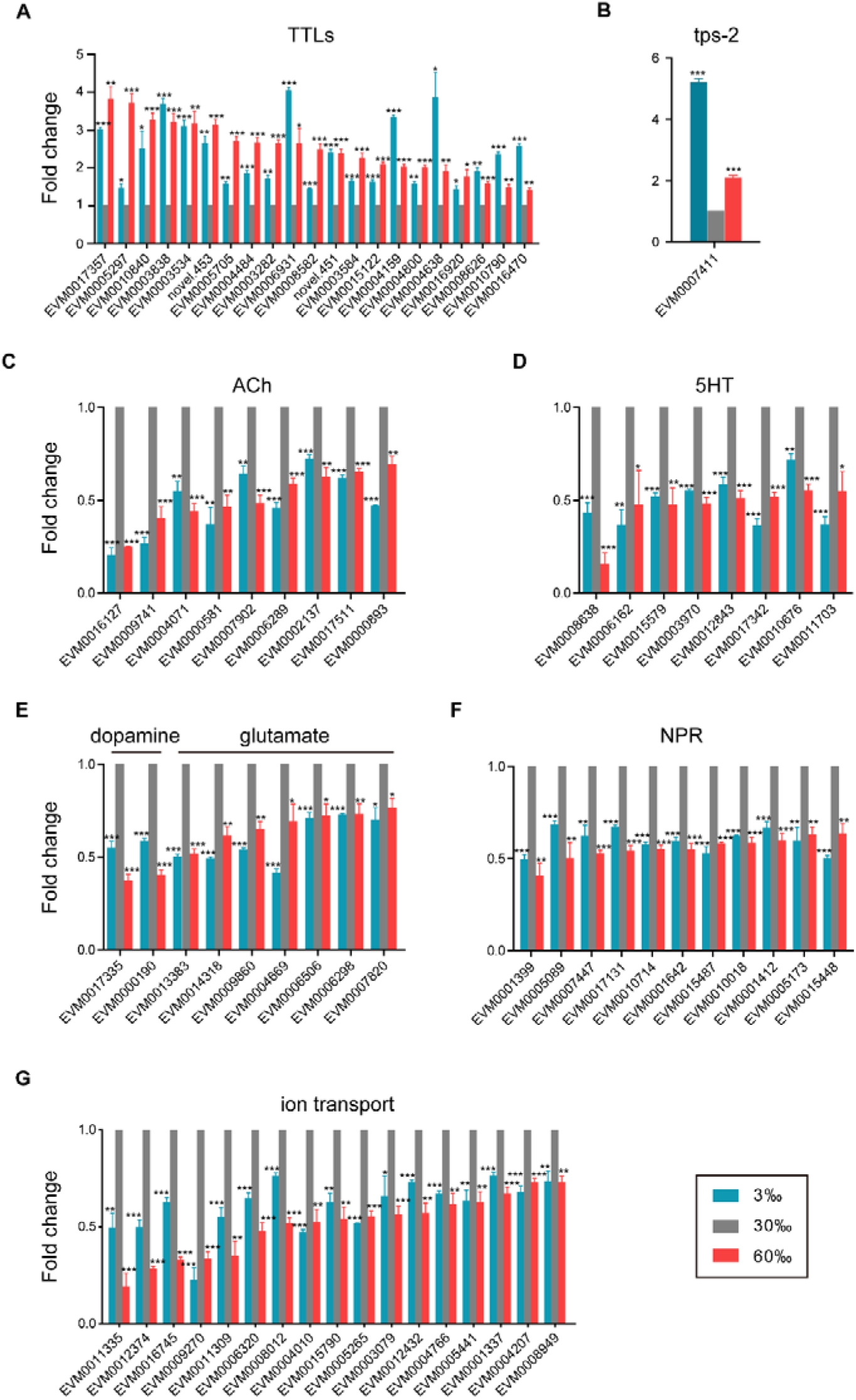
Shared transcriptomic signature under both hyposaline and hypersaline environments. **(A)** Expression level of transthyretin-like family genes (TTLs). **(B)** Expression level of trehalose-6-phosphate synthase gene (tps-2). **(C)** Expression level of acetylcholine receptor genes (ACh). **(D)** Expression level of serotonin receptor genes (5HT). **(E)** Expression level of dopamine and glutamate receptor genes. **(F)** Expression level of neuropeptide receptor genes (NPR). **(G)** Expression level of ion transporter genes. Fold change indicates the ratio of the treatment group (3‰, 60‰, as indicated) to the control group (30‰). The error bars represent standard error of the mean of three biological replicates per condition. \**P* < 0.05, \*\**P* < 0.01, \*\*\**P* < 0.001.

In addition, the trehalose-6-phosphate synthase gene (EVM0007411/*tps-2*, **Figure 3B**), which is crucial for trehalose biosynthesis, was significantly induced upon both salinity stresses.

Moreover, we found that multiple neurotransmitter receptor genes were down-regulated in both conditions (**Figure 3C-F**). The expression levels of seven nicotinic acetylcholine receptor genes (EVM0016127/*eat-2*, EVM0009741/*eat-2*, EVM0006289/*acr-3*, EVM0007902/*acr-5*, EVM0000893/*acr-11*, EVM0017511/*acr-12* and EVM0000581), as well as two muscarinic acetylcholine receptor genes (EVM0002137/*gar-2* and EVM0004071), were significantly down-regulated in both hyposaline and hypersaline conditions (**Figure 3C**). Similarly, eight serotonin receptor genes (such as EVM0012843/*ser-1*, EVM0010676/*ser-2* and EVM0015579/*ser-7*, **Figure 3D**), two dopamine receptor genes (EVM0000190/*dop-1* and EVM0017335, **Figure 3E**), seven glutamate receptor genes (EVM0013383/*glc-4*, EVM0014318/*glc-2*, EVM0009860/*mgl-1*, EVM0004669/*avr-15*, EVM0006506/*ggr-2*, EVM0006298/*avr-14* and EVM0007820/*glr-1*, **Figure 3E**), and eleven neuropeptide receptor genes (such as EVM0010018/*npr-4*, EVM0015448/*npr-6*, EVM0005173/*npr-15*, EVM0001642/*ckr-1*, EVM0017131/*ckr-2*, EVM0007447/*frpr-9*, EVM0001412/*lat-2* and EVM0010714/*lat-2*, **Figure 3F**) were all significantly down-regulated. These results implied that certain neuronal related signaling transduction processes were severely impaired by short-time stresses caused by both low and high salinity.

Additionally, based on studies on fishes and marine invertebrates, ion transporters and channels are key components of osmoregulation (Niu et al., 2020;Vij et al., 2020;Zhang et al., 2020a). In the present study, a dozen of V-type H^+^-transporting ATPase genes (EVM0006836/*vha-1*, EVM0002567/*vha-3*, EVM0005735/*vha-4*, EVM0007934/*vha-5*, EVM0008894/*vha-5*, EVM0000966/*vha-7*, EVM0001861/*vha-8*, EVM0014072/*vha-12*, EVM0014618/*vha-13*, EVM0014647/*vha-15*, EVM0015095/*vha-16* and EVM0006795/*vha-19*) were enriched and showed elevated expression under hyposaline condition (**Supplementary Figure 1**). The upregulation of these genes was also reported in other marine invertebrates, including the mud crab *Scylla paramamosain* (Niu et al., 2020) and the shrimp *Litopenaeus vannamei* (Wang et al., 2012), indicating their conserved function in response to low salinity stress among marine invertebrates. However, most of those V-type H^+^-transporting ATPase genes were also upregulated under hypersaline condition in *L. marina* (**Supplementary Figure 1**), suggesting a specific role in marine nematodes. On the other hand, a battery of ion channel and transporter genes such as potassium channel genes (EVM0012374/*twk-24*, EVM0015790/*shw-3*, EVM0004766/*kcnl-3* and EVM0004207/*kcnl-2*), sodium channel genes (EVM0009270/*egas-2* and EVM0008949/*nhx-8*), cyclic nucleotide gated channel gene (EVM0006320/*tax-4*), potassium/chloride transporter gene (EVM0004010/*kcc-2*), transient receptor potential cation channel genes (EVM0012432/*trp-1*, EVM0001337/*trp-2* and EVM0005441/*osm-9*) were identified, demonstrating down-regulation in both stress environments (**Figure 3G**), reflecting their association with the ionic homeostasis under salinity stresses.

Overall, these shared features indicate the existence of conserved strategies for response to stressful salinity environments in *L. marina*.

### 3.4 Up-regulated Genes under Low Salinity Condition

Hyposaline (3‰) impacted *L. marina* development, we thus further analyzed genes that were induced specifically under this low salinity condition, and detected 144 DEGs (**Figure 4A**). For instance, fatty acid desaturase genes (EVM0008235/*fat-2*, EVM0001302/*fat-4* and EVM0011847/*fat-3*), very long chain fatty acid elongase genes (EVM0013022/*elo-2*, EVM0000114/*elo-6* and EVM0000630/*elo-5*), very-long-chain enoyl-CoA reductase gene (EVM0000944/*art-1*), and long-chain-fatty-acyl-CoA reductase gene (EVM0015936/*let-767*), involved in biosynthesis of unsaturated fatty acids (UFAs), were significantly accelerated (**Figure 4B**). These results suggested that UFAs might play important roles in *L. marina*’s responding to low salinity stress.

**FIGURE 4.**
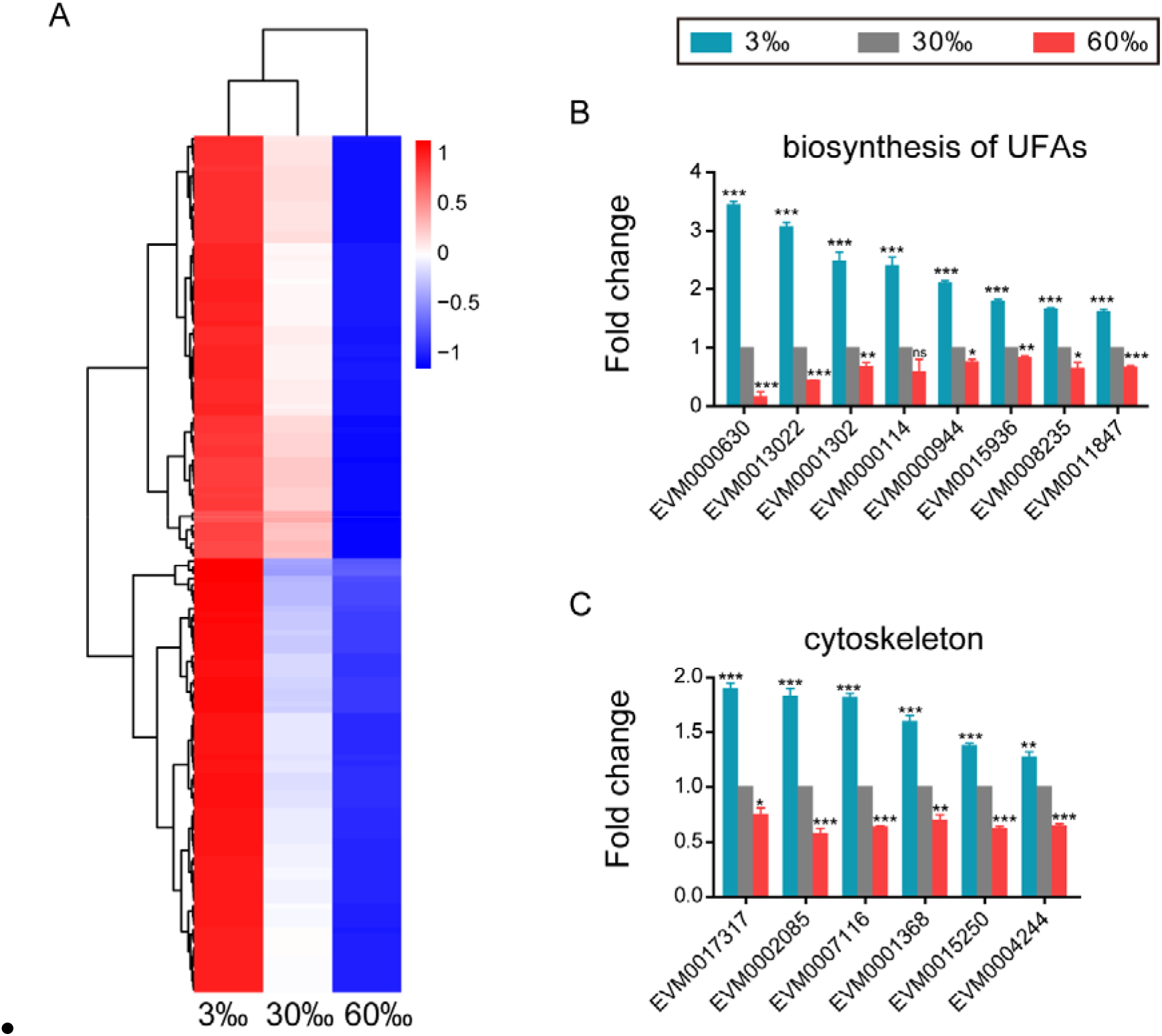
Genes that up-regulated in expression with decreasing salinity. **(A)** Heatmap of DEGs that up-regulated in expression with decreasing salinity. The scale bar shows the z-score for a differentially expressed gene. Red indicates upregulation; blue indicates downregulation. **(B)** Expression level of genes involved in biosynthesis of unsaturated fatty acids (UFAs). **(C)** Expression level of cytoskeleton tubulin and related intraflagellar transport (IFT) genes. Fold change indicates the ratio of the treatment group (3‰, 60‰, as indicated) to the control group (30‰). The error bars represent standard error of the mean of three biological replicates per condition. \**P* < 0.05, \*\**P* < 0.01, \*\*\**P* < 0.001.

In addition, we observed specifically up-regulation in four tubulin genes (EVM0017317/*tba-4*, EVM0007116/*tba-5*, EVM0015250 and EVM0004244/*ben-1*, **Figure 4C**) and two intraflagellar transport (IFT) genes (EVM0001368/*osm-3* and EVM0002085/*daf-10*, **Figure 4C**) under low salinity condition.

However, the above UFAs, tubulin and IFT genes showed significantly opposing changes between low and high salinity stresses (**Figure 4B-C**), indicating their critical roles in salinity stress response.

### 3.5 Up-regulated Genes under High Salinity Condition

Although worms hardly survived under high salinity stress, 192 DEGs were found having the highest expression levels under this extreme condition (**Figure 5A**). Dozens of cuticle related collagen genes were up-regulated under 60‰ salinity, including EVM0002243/*col-156*, EVM0010502/*dpy-5*, EVM0000427/*col-77*, EVM0006032/*col-86*, EVM0016231/*col-166*, EVM0005554/*col-107*, EVM0001263/*col-104*, EVM0001382/*lon-3*, EVM0003934/*col-149*, EVM0008240/*dpy-17*, EVM0011594/*sqt-3*, and EVM0000108/*col-93* (**Figure 5B**). It is likely reflecting essential roles of these collagen genes in response to high salinity. Of note, most of the above collagen genes showed significantly opposing changes under low and high salinity stresses (**Figure 5B**).

**FIGURE 5.**
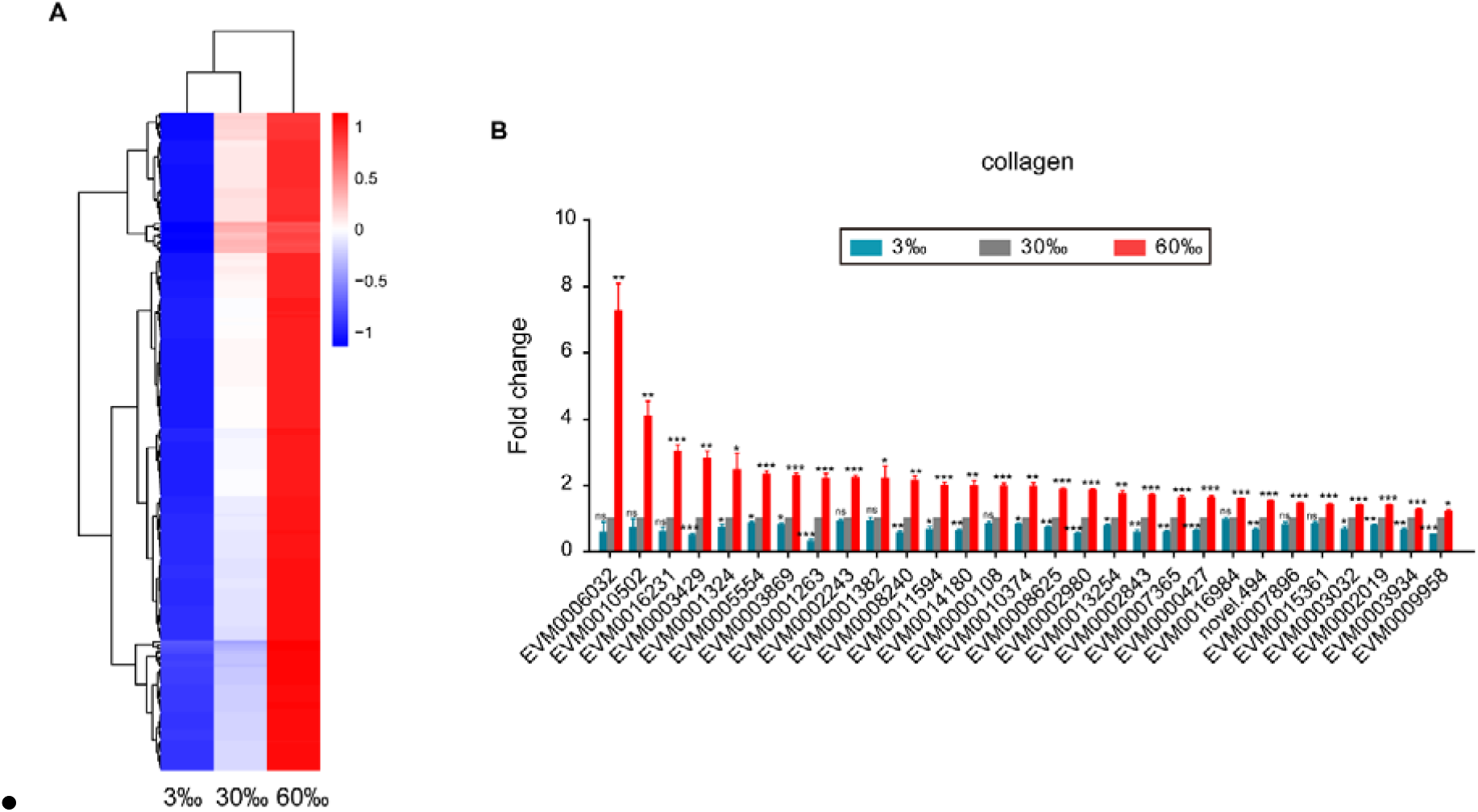
Genes that up-regulated in expression with increasing salinity. **(A)** Heatmap of DEGs that up-regulated in expression with increasing salinity. The scale bar shows the z-score for a differentially expressed gene. Red indicates upregulation; blue indicates downregulation. **(B)** Expression level of cuticle collagen genes. Fold change indicates the ratio of the treatment group (3‰, 60‰, as indicated) to the control group (30‰). The error bars represent standard error of the mean of three biological replicates per condition. \**P* < 0.05, \*\**P* < 0.01, \*\*\**P* < 0.001.

### 3.6 Quantitative Real-time PCR Validation

We applied qPCR to validate the expression patterns of interest genes identified from our RNA-seq results. Consistent trends were detected and shown as in Figure 6. The expression levels of transthyretin-like family gene EVM0003534, trehalose-6-phosphate synthase gene EVM0007411/*tps-2*, were significantly increased under both 3‰ and 60‰ salinity conditions (**Figure 6**). While, the expression levels of dopamine receptor gene EVM0000190/*dop-1*, glutamate receptor gene EVM0013383/*glc-4*, acetylcholine receptor gene EVM0009741/*eat-2*, serotonin receptor gene EVM0012843/*ser-1*, neuropeptide Y receptor genes EVM0015448/*npr-6* and EVM0010018/*npr-4*, ion transporter genes EVM0004010/*kcc-2* and EVM0012374/*twk-24*, were significantly decreased under both conditions (**Figure 6**). In addition, we confirmed that fatty acid elongase gene EVM0013022/*elo-2*, fatty acid desaturase gene EVM0001302/*fat-4*, tubulin gene EVM0007116/*tba-5*, were up-regulated under hyposaline whereas down-regulated under hypersaline environment (**Figure 6**). By contrast, the expression of cuticle collagen genes EVM0002243/*col-156* and EVM0005554/*col-107* were validated showing specific upregulation under high salinity stress (**Figure 6**).

**FIGURE 6.**
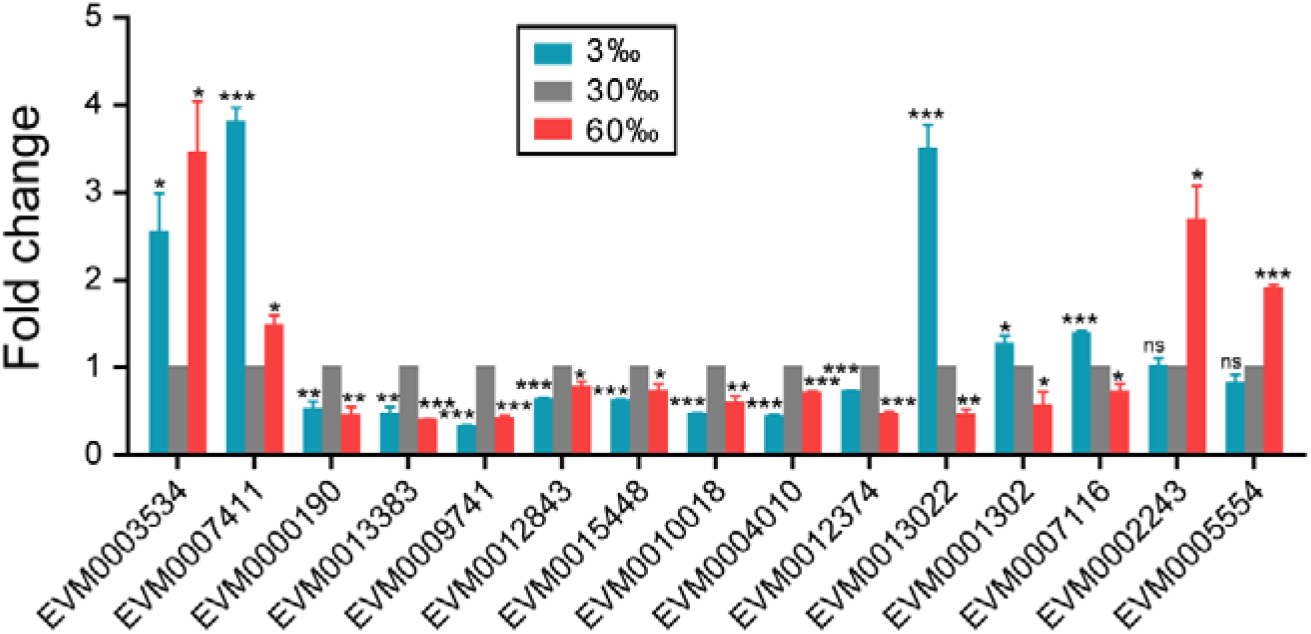
Validation of the RNA-seq results using qPCR. Fold change indicates the ratio of the treatment group (3‰, 60‰, as indicated) to the control group (30‰). The mean fold changes and standard error of the mean of three biological replicates are graphed. \**P* < 0.05, \*\**P* < 0.01, \*\*\**P* < 0.001.

## 4 Discussion

### 4.1 Synchronization in *L. marina* for Large-Scaled Analysis

According to the synchronization methods used for hermaphroditic worms such as *C. elegans* and *P. pacificus* (Pires da Silva, 2005), we could not obtain enough L1 larvae samples for large-scale analysis in terms of the dioecious marine nematode *L. marina*. Therefore, we instead allowed *L. marina* adults to lay eggs on plates overnight, and then collected eggs from these plates. After a short-time treatment with bleaching solution to kill *E. coli* OP50 in the washing solution, eggs were subsequently incubated in filtered sterile seawater to hatch. Finally, synchronized newly hatched *L. marina* L1 larvae were effectively obtained by filtration using a grid nylon filter with mesh size of 25 μm. The establishment of large-scale L1 larvae collecting protocol developed in this study will facilitate further *L. marina* multi-omics studies, which requires large-scale synchronized worms.

### 4.2 Marine Nematode *L. marina* Has a Wider Range of Salinity Tolerance

In the intertidal areas, habitat salinity of *L. marina* is subject to either sudden or gradual changes in response to tides, rainfall, ocean currents, seawater evaporation and climate. In the laboratory, we found that *L. marina* could survive a wider salinity from 3‰ to 60‰. Of note, for marine nematodes, 60‰ salinity is obviously an extreme condition, which is almost twice of that of sea water. By contrast, we noticed that *C. elegans* couldn’t survive at 30‰ salinity (30 L1s per plate in triplicates). Thus, *L. marina* is a euryhaline marine nematode and has a wider range of salinity tolerance than terrestrial nematode *C. elegans*. Further studies using marine nematode *L. marina* as a model, will provide universal mechanisms underlying marine invertebrates’ euryhaline adaptation.

### 4.3 Transthyretin-Like (TTL) Family Genes are Presumably Involved in the Damage Control Mechanisms in Response to Salinity Stresses

Under diverse environmental and physiological stresses, organisms usually demonstrate various degrees of cell damage by stress-induced protein misfolding, denaturation or aggregation, thereby disrupting proteostasis and cell homeostasis (Lamitina et al., 2006;Galluzzi et al., 2018). In the process of stress response, common stress-inducible genes, such as heat shock protein genes, are induced to protect cells (Spees et al., 2002;Lamitina et al., 2006). Such stress-inducible genes were also observed in our transcriptome results, for example, a series of HSP20, HSP70 family chaperone genes and dozens of proteasome related genes were significantly up-regulated under both low and high salinity stresses (**Supplementary file 3**).

In terms of shared common DEGs between both salinity stresses, one prominent type of significantly up-regulated were the transthyretin-like (TTL) family genes. In *L. marina*, at least 38 TTLs family genes have been annotated by database mining. In the present study, a total of 21 genes encoding TTLs were up-regulated under both hyposaline and hypersaline environments, suggesting that they might play important roles in responding to salinity stresses in *L. marina*. TTLs represent one of the largest nematode-specific protein families, sharing sequence similarity to vertebrate transthyretins (Parkinson et al., 2004). In vertebrates, transthyretins are present in extracellular fluids to transport thyroid hormones as well as vitamin A (Vieira and Saraiva, 2014). In terms of nematode TTLs, they were presumed to participate in disposal of toxic lipophilic moieties and hormonal signaling (Parkinson et al., 2004;Jacob et al., 2007). However, these elusive genes have not been implicated in the response to salinity stress up to now and their functions are largely unknown. TTR-52 was reported as a bridging factor involved in cell corps engulfment and apoptosis (Wang et al., 2010;Mapes et al., 2012), indicating that up-regulation of TTL genes might be part of the damage control mechanisms in response to either low or high salinity stresses in *L. marina*.

### 4.4 Trehalose-6-phosphate synthase gene (TPS) is up-regulated in *L. marina* upon both Low and High Salinity Stresses

Cells of almost all organisms accumulate organic osmolytes when exposed to hyperosmolarity, and more than one type of osmolytes can be utilized for a particular organism (Burg and Ferraris, 2008). Unlike most marine invertebrates, which mainly use free amino acids and methylamines as organic osmolytes (Niu et al., 2020), it has well demonstrated that hyperosmotic stress in *C. elegans* activates rapid accumulation of glycerol via the rapid up-regulation of the glycerol-3-phosphate dehydrogenate enzyme *gpdh-1*, a key gene for de novo glycerol synthesis, thereby to balance the osmotic pressure (Lamitina et al., 2004;Lamitina et al., 2006). Based on the annotation information of *L. marina* genome, the predicted glycerol-3-phosphate dehydrogenase gene, EVM0001663/*gpdh-1*, was shown responsive to hypersaline stress in the current study, which was significantly up-regulated under high salinity (**Supplementary Figure 2**), indicating that *L. marina* might utilize glycerol as an osmolyte in response to high salinity stress similar to *C. elegans*.

Trehalose, a disaccharide of glucose, is present in a wide variety of organisms including nematodes, and is known to act as stress protectant to against effects of dehydration, desiccation, heat, freezing as well as high osmotic stress (Wharton, 2003;Erkut et al., 2011;Hibshman et al., 2020). It not only supports survival by stabilizing lipid membranes and improving proteostasis during water loss, but also serves as an energy source. It is known that trehalose-6-phosphate synthase gene (TPS) encodes the enzyme catalyzing the first step of trehalose biosynthesis (Watts and Ristow, 2017). Previously, it was reported that trehalose levels were elevated and conferred hypertonic stress resistance in *C. elegans age-1* mutants, which was suppressed by RNAi knockdown of TPS genes, indicating an important functional role of TPS in hypertonic stress resistance in *C. elegans* (Lamitina and Strange, 2005). In this study, we observed that *L. marina* TPS gene, EVM0007411/*tps-2*, was significantly up-regulated under both hyposaline and hypersaline stresses (**Figure 3B**), which could possibly cause accumulation of trehalose to facilitate its adaptation to both stress environments.

### 4.5 Certain Neuronal Signaling are Transcriptionally Repressed by Salinity Stresses

In *C. elegans*, acetylcholine, serotonin, dopamine, glutamate and neuropeptide are known important neurotransmitters, which have been well demonstrated their involvement in a broad repertoire of behaviors, including locomotion, feeding, reproduction, social behavior, mechanosensation, chemosensation, learning, memory, behavioral plasticity and adaptation (Chase and Koelle, 2007;Rand, 2007;Hukema et al., 2008;Frooninckx et al., 2012;Vidal-Gadea and Pierce-Shimomura, 2012). Under harsh conditions, the nervous system plays critical roles in worm stress response, facilitating worm survival and adaptation (Kim and Jin, 2015). Although, specific neurons that function in osmotic stress response have studied extensively, for example, it is well known that ASH neurons are required for hyperosmotic sensation and ASEs are for hypoosmolarity, both types of neurons function in *C. elegans* osmotic avoidance behavior, allow worms to respond rapidly to escape from harmful osmotic stress conditions (Hilliard et al., 2005;Kunitomo et al., 2013). When osmotic stress is unavoidable, worms engage a sleep-like quiescent behavior and cease locomotion and feeding, which is dependent on ALA neuron (Hill et al., 2014). However, specific neurotransmitters, including serotonin, dopamine, glutamate and neuropeptide are only reported affecting avoidance behavior to NaCl in *C. elegans* (Hukema et al., 2008;Watteyne et al., 2020). To our knowledge, this is the first to describe a systematic repressed cholinergic, serotonergic, dopaminergic, glutamatergic and neuropeptide signaling in response to both low and high salinity stresses in *L. marina*.

In the present study, these reduced neural signaling are positively correlated with the reduced worm mobility in salinity stress environments. Furthermore, we speculate that these reduced signaling might further activate osmoregulation pathways in *L. marina* to promote its adaptation to the stressed salinity conditions.

### 4.6 Unsaturated Fatty Acids (UFAs) are Involved in Hyposaline Stress Response in *L. marina*

Genes involved in UFAs biosynthesis were observed to be significantly up-regulated in low salinity condition. Defects in UFAs biosynthesis have been reported to cause deficiencies in worm growth, development and neurological function (Watts and Browse, 2002;Kniazeva et al., 2003). Notably, we found that several genes encoding fatty acid elongases (EVM0013022/*elo-2*, EVM0000630/*elo-5*) and fatty acid desaturases (EVM0008235/*fat-2*, EVM0011847/*fat-3*) were significantly decreased under hypersaline stress, which might account for the developmental defects in 60‰ salinity stressed worms.

UFAs have a profound effect on the fluidity, flexibility and permeability of cell membranes, as well as play important roles in energy storage and signaling process (Zhu and Han, 2014). The synthesis of UFAs is regulated during changing environmental conditions, playing a crucial role in environmental adaptation mechanism in organisms, such as animals, plants and microorganisms (Wada et al., 1990;Miquel et al., 1993;Svensk et al., 2013;Yancey, 2020;Zhang et al., 2020b). It has been widely reported that fatty acids were more unsaturated in salt-tolerant plants, yeast as well as bacterial cells (de Carvalho et al., 2014;Guo et al., 2019;Li et al., 2019). Moreover, Lucu *et al.* demonstrated that UFAs might be important during the acclimation of the shore crab *Carcinus aestuarii* to hypoosmotic condition (Lucu et al., 2008). However, nematode UFAs have not been implicated in the response to salinity stress as far as we know. Under hyposaline stress, body swelling is initially observed for *L. marina* L1s, due to the influx of fluid. While worms did present a better ability to tolerate and acclimate to this condition, showing relatively normal development and fertility afterwards. It suggests that the regulatory mechanism of body volume recovery is more effective under hyposaline stress. This may be related to the cell membrane properties and functions in *L. marina*. Thus, we propose that the effective induction of UFAs biosynthesis genes may acts as part of the protective and adaptative strategies of marine nematodes upon low salinity stress.

### 4.7 Cuticle Collagen Genes are Involved in Hypersaline Stress Response in *L. marina*

Collagens are major structural proteins for the nematode exoskeleton, cuticle. The identified *C. elegans* cuticle collagen mutants usually show either body morphology defects or locomotion defects, which is consistent with the cuticle’s essential role for maintenance of body shape, as well as movement via attachments to muscles (Page and Johnstone, 2007). Moreover, several cuticle collagen mutants, such as *dpy-2*, *dpy-7* and *dpy-10*, were reported to exhibit constitutive activation of *gpdh-1* expression and glycerol accumulation, and show osmotic resistance phenotype (Lamitina et al., 2006;Wheeler and Thomas, 2006;Dodd et al., 2018). It is believed that the cuticle may serve as a sensor for osmotic stress and play important roles in *C. elegans* osmotic regulation (Lamitina et al., 2006;Dodd et al., 2018).

The cuticle in *C. elegans* is synthesized and secreted by underlying hypodermis, this process occurs five times during development, first at the end of embryogenesis before hatching and then again at the end of each larval stage before molting. *C. elegans* cuticle collagens are encoded by a multi-gene family, consisting over 170 genes (Page and Johnstone, 2007). In fact, these genes are not all expressed at the same time during cuticle synthesis. Clear spatial and temporal differences can be observed for individual genes (Johnstone, 2000;Page and Johnstone, 2007), indicating each cuticle collagen gene has specific roles in cuticle formation and function. The cuticle functions as the primary barrier between worm and its environment, and acts as the first line of defense against environmental stresses (Page and Johnstone, 2007). Previously, Dodd *et al.* has shown that multiple cuticle collagen genes can be induced when *C. elegans* exposed to high NaCl (Dodd et al., 2018). Recently, we reported that a battery of collagen genes in *C. elegans* increased their expression to deal with acidic pH stress environments (Cong et al., 2020). Together, these results indicated a protective role of collagens in response to various stresses. Similarly, in the present study, an abundant group of over 20 cuticle collagen genes were significantly up-regulated upon hypersaline stress, while most of them were remarkably down-regulated upon hyposaline stress, indicating certain collagens could specifically function to detect or transform salinity stress-induced signals, or just change the chemical and physical composition of the cuticle to provide the primary barrier to defend the dynamic osmotic variation.

### 4.8 Cytoskeleton related genes are Differentially Regulated under Different Salinity Stresses in *L. marina*

Tubulin is the basic component of cytoskeleton microtubules. It plays an indispensable role in structure maintaining, neuronal sensation as well as many other cell processes including intraflagellar transport (IFT). In our results, four tubulin genes, such as EVM0017317/*tba-4*, EVM0007116/*tba-5*, EVM0015250 and EVM0004244/*ben-1*, exhibited specific up-regulation when salinity is decreasing (**Figure 4C**). The *tba-4* gene encodes α-tubulin in *C. elegans*, both TBA-5 and BEN-1 are neuronal tubulins (Hurd, 2018). TBA-5 is an axonemal α-tubulin expressed in amphid and phasmid sensory neurons where it is localized to cilia. BEN-1 is a neuronal β-tubulin in *C. elegans*, which is also broadly expressed in the nervous system. Moreover, it is reported that IFT is essential for assembly, maintenance and function of sensory cilia in *C. elegans* (Hao et al., 2011). Here, we found that two IFT genes (EVM0001368/*osm-3* and EVM0002085//*daf-10*) were induced expression specifically under low salinity condition (**Figure 4C**). The *osm-3* gene in *C. elegans* encodes a kinesin-2 family member of IFT motors, mediating IFT particles transport within sensory cilia (Prevo et al., 2015). *daf-10* is required for IFT and for proper development of a number of sensory neurons (Bell et al., 2006). By contrast, both *tba-5* and *daf-10* genes were found to be upregulated in the osmo-resistant *dpy-7* worms (Dodd et al., 2018). Moreover, the above tubulin and IFT genes exhibited significantly opposing changes between low and high salinity stresses (**Figure 4B-C**), indicating their critical roles in salinity stress response.

In conclusion, we have described for the first time the genome-wide transcriptional responses to both hyposaline and hypersaline stresses in the marine nematode *L. marina*. The present study will provide an essential foundation for identifying the key genes and genetic pathways required for osmoregulation in the marine nematodes. Given a wide range of salinity tolerance of the marine nematodes, our results and further genetic analysis of key gene(s) of osmoregulation in *L. marina* will likely provide important insights into biological evolution and physiological adaptation mechanisms in nematodes and other organisms in general.

## 5 Article types

Original Research Article.

## 7 Author Contributions

YX and LZ conceived and designed the experiments. YX carried out most of the experiments, analyzed the data, and wrote the manuscript. PZ contributed to the RNA-seq sampling and qPCR validation. LZ edited the manuscript and supervised the project. All authors read and approved the final manuscript.

## 9 Acknowledgments

We are grateful to all members of the LZ laboratory for their helpful discussions.

## 10 Data Availability Statement

The datasets generated for this study can be found in NCBI, and the BioProject ID: PRJNA694479.

## 11 Supplementary Figures

**Supplementary FIGURE 1.**
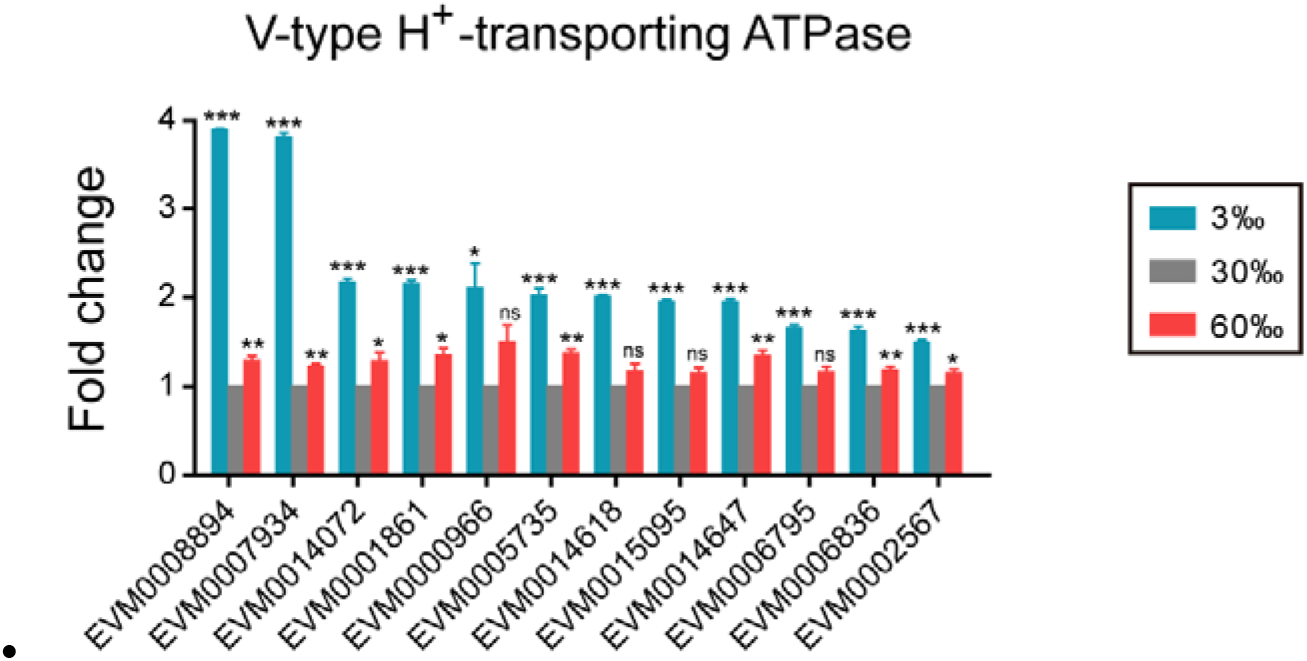
Expression level of V-type H+-transporting ATPase genes upon salinity stresses in *L. marina*. Fold change indicates the ratio of the treatment group (3‰, 60‰, as indicated) to the control group (30‰). The error bars represent standard error of the mean of three biological replicates per condition. \**P* < 0.05, \*\**P* < 0.01, \*\*\**P* < 0.001.

**Supplementary FIGURE 2.**
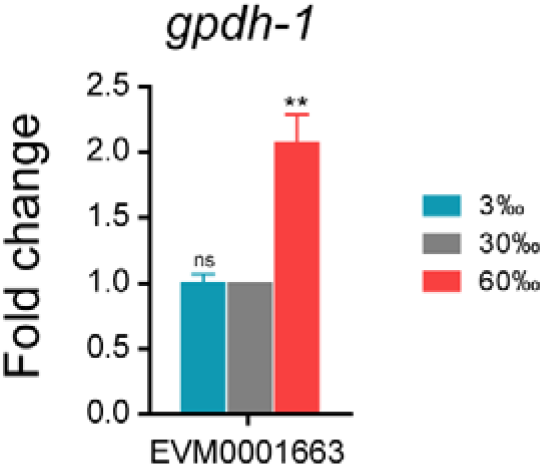
Expression level of *L. marina* glycerol-3-phosphate dehydrogenase gene upon salinity stresses. Fold change indicates the ratio of the treatment group (3‰, 60‰, as indicated) to the control group (30‰). The error bars represent standard error of the mean of three biological replicates per condition. \**P* < 0.05, \*\**P* < 0.01, \*\*\**P* < 0.001.

